# Early *Candida*–Oral Tumor Interactions Suggest miRNA-Mediated Regulation of Inflammatory and Tumor-associated processes

**DOI:** 10.1101/2025.06.05.657835

**Authors:** Renata Toth, Marton Horvath, Julian R. Naglik, Attila Gacser

## Abstract

When simultaneously present, oral cancer and oral candidiasis are associated with high mortality. Recently, fungi-driven OSCC has emerged as a recognized phenomenon, that is triggered by *Candida* and sustained by several pro-tumor mechanisms. This study aimed to deepen our understanding and investigate how these mechanisms are regulated at the early stages of fungi-tumor interactions. Post-transcriptional regulation was examined through the analysis of miRNA (miR) responses in a metastatic cell line. We characterized early tumor responses to two common oral yeasts, *C. albicans*, frequently responsible for oral infections, and *C. parapsilosis*, a yeast with mostly commensal behavior in this niche. Exposed to *C. albicans*, tumor cells exhibited overexpression of inflammatory and pro-tumor pathways. This was also mirrored at the miR-level. Intriguingly, this species induced upregulation of oncogenic miRs, while simultaneously downregulated tumor-suppressive miRs. miR-target analyses revealed targets primarily to be involved in inflammation (TNFa signaling via NFKB), albeit tumor progression was also heavily influenced based on the targets identified within the PanCancer Progression Panel. Several miR-target pairs were identified with TRIB1-miR-30c-1-3p highlighted at the route of inflammation, and MMP3-miR-374a-3p and JUN-miR-1268b/miR-16-1-3p/miR-30c-1-3p at the route of OSCC progression. These findings highlight that *C. albicans* drives OSCC progression, in part, through miR regulation. Contrarily, *C. parapsilosis* elicited tumor responses predominantly related to hypoxia and metabolic regulation, followed by a mild activation of inflammatory and tumorigenic processes. No significant miR-regulatory responses were identified. To sum up, we added a new layer to fungi-tumor interactions and identified regulatory elements, that might hold future translational potential.

**Importance:** Previous studies have shown that yeast carriage in the oral cavity of oral cancer patients is significantly higher than those of healthy individuals. Additionally, the abundance of fungal cells is higher at malignant epithelial surfaces compared to the unaffected mucosal tissues. These observations indicated a direct interaction between mouth residing yeasts and oral tumorous cells. We have previously demonstrated that certain *Candida* species actively contribute to tumor progression in metastatic oral epithelial cells via various mechanisms including the promotion of migration, MMP activity or oncometabolite secretion. In this study, our findings demonstrate, that fungi-driven OSCC progression is also evident at the level of miRNA regulation, when *C. albicans* is present. In this context, the fungus might also enhance existing post-transcriptional regulatory mechanisms in the malignant oral epithelium. This behavior seems to be species-specific, as another common resident of the oral microflora, *C. parapsilosis* does not trigger such an effect.

## Introduction

In recent decades, the human microbiome has been frequently associated with cancer. While the exact cause-and-effect relationship remains unclear, alterations in the niche-specific microbiota can still serve as a diagnostic marker for various types of cancer (1). In certain cases, specific microbes within this altered niche can be identified and directly linked to the development of a disease. Viruses such as HPV (cervical and oropharyngeal cancers) and EBV (Burkitt lymphoma and Hodgkin lymphoma), as well as bacteria like *Helicobacter pylori* (gastric cancer), are well-documented direct oncogenic factors, and are classified as Group I carcinogens by the IARC (2). However, fungal species have yet to be conclusively identified as a direct driver of cancer. Although not listed as carcinogens, fungi such as *Aspergillus* spp., *Fusarium* spp., and *Candida* spp. might also contribute to pro-tumor processes, potentially through inducing chronic inflammation or due to the carcinogenic effects of the toxins they produce (3).

In our previous study, we revealed that *Candida albicans* in fact acts as a direct driver of tumor progression, particularly in oral cancer, and also contributes to carcinogenesis (4). The study highlighted that *C. albicans* provokes oral squamous cell carcinoma (OSCC) progression by enhancing pro-tumor mechanisms, such as matrix metalloprotease (MMP) activity, tumor cell migration, and the overrepresentation of metastatic genes and markers of epithelial-to-mesenchymal transition (EMT). Mild to moderate dysplasia was also observed in *C. albicans*-colonized mice tongues, supporting the idea that this species also contributes to oncogenesis under predisposing conditions. Subsequent studies have also confirmed the association between *C. albicans* presence and oral malformations (5, 6), and revealed the *C. albicans*-driven upregulation of programmed death ligand-1 (PD-L1) (7) and the dysregulation of the KRAS signaling pathway and E2F target downstream genes (8). While light has been shed on certain biological processes driving the observed phenomena, still much remains to be understood. For instance, how the disrupted cellular functions are regulated. Identifying key regulators could open new avenues for intervention.

Small non-coding RNAs, specifically microRNAs (miRs), are crucial regulators of cellular responses, controlling nearly all signaling pathways, from cell cycle regulation to stress and inflammatory responses (9). As numerous studies have implicated the broad applicability of miRs (10–12), and a Nobel Prize has also recently been awarded to the field (13), miRs are suggested to hold great potential and a promising role in future medicine.

Investigating the role of miRs in fungi-driven tumor progression could not only add a new dimension to fungi-tumor interactions but also offer a new perspective on potential interventions. Such insights could lead to novel opportunities in either early detection (diagnostic potential) or therapeutic interventions.

Our previous research provided valuable insights into how fungal presence regulates host responses in oral epithelial cells with a healthy origin (14). During the study, major transcriptome and miRome changes were identified after *Candida* stimuli. Alterations in both mRNA and miR profiles revealed that oral epithelial cells actively discriminate between *Candida albicans* and *Candida parapsilosis* based on their pathogenic potential, and the host responses varied by species, dose, and time. Our findings revealed an altered, miR-based regulation of inflammatory-, hypoxia- and carbohydrate metabolic pathways. Albeit interestingly, under specific conditions certain tumor-associated pathways were also dysregulated. These results suggested early signs for *Candida* species actively influencing tumorous processes.

Given the long-standing association between oral cancer and *Candida* species (15–25), and the recent evidence for direct interactions between *C. albicans* and oral tumor cells, the aim of this study was to identify miR-associated regulatory mechanisms during the early stages of fungi-tumor interactions. With this work, we seek to add a new layer to fungi-tumor interactions and to identify regulatory elements, that might offer a new perspective on potential interventions.

In this study, we aimed to test the early regulatory effect of *C. albicans*—the most frequently identified fungal species in oral tumors—and *C. parapsilosis*, - another common resident of the oral mycobiome, though typically more commensal than pathogenic - on metastatic OSCC cells.

## Results

### *Candida*-driven tumor progression: species-, dose- and time-dependence in the metastatic cell line

Microbes can contribute to tumor progression either via inflammatory or non-inflammatory processes.

Fungi-associated tumor progression is specifically linked to either enhanced inflammation or to the overactivation of protumor markers. Thus, we aimed to investigate both events. Furthermore, we aimed to address the question whether the *Candida*-induced oral tumor progression effect is dose- or time-dependent. For these studies, we applied two distinct species, *C. albicans* - the primary species associated with oral cancer - and *C. parapsilosis* - a less frequent pathogen, rather, a commensal-like species of the oral mycobiome. We examined their effect on the metastatic cell line, HSC-2, in various doses.

As a result, we found that *C. albicans* promotes both microbe-induced tumor progression routes, thus contributes to tumor progression both indirectly (via inflammation) and directly (via tumor marker genes). As shown on Figure 1, the secretion of major inflammatory cytokines (IL-1b, IL-6 and IL-8, except for CCL5; after 24 hours) increased substantially (FIG1A), and several OSCC-tumor marker genes (JUN1, LAMC2, MMP1, MMP10 and INHBA; after 6 hours) also showed a markedly increased expression in its presence (FIG1B). Interestingly, there was no significant difference in the secretion of CCL-5 compared to the control. When further investigating the effects, we observed that the increase in fungal load directly correlated with the elicited cytokine response, but not with the expression of tumor-marker genes. Increase in the expression of oncogenes required a couple of hours of fungal stimuli (6 hours), as such differences did not seem to manifest as early as 1 hour of the interaction (FIG S1).

**Figure 1.**
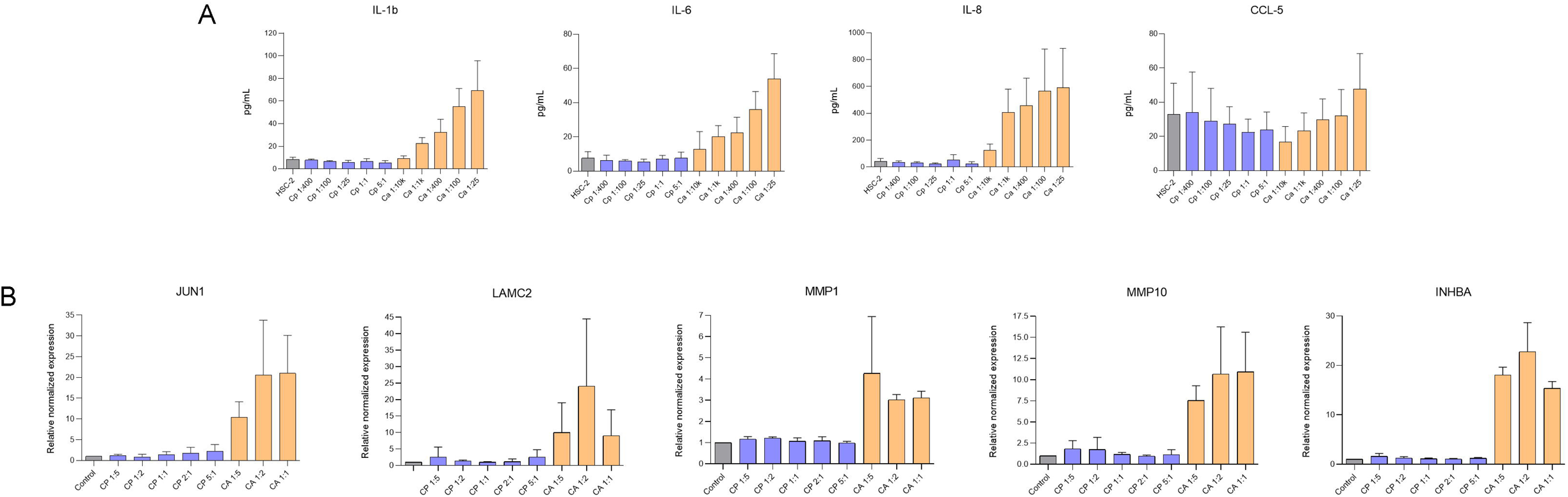
HSC-2 epithelial cell responses to *C. albicans* and *C. parapsilosis.* (A) Secreted inflammatory cytokines (pg/mL) by HSC-2 cells 24 hours after *Candida* exposure, at the shown MOIs (pathogen:host). (B) Relative normalized expression of pro-tumor marker genes in HSC-2 cells, 6 hours after *Candida* stimuli, at different infection doses (pathogen:host). Uninfected HSC-2 cells served as controls.

Contrarily to *C. albicans*, *C. parapsilosis* failed to elicit any of the above-mentioned responses, as all triggered responses were equivalent to that of the untreated control condition, regardless of the fungal load, or the length of the stimulus (FIG1, FIG S1).

### Early transcriptional responses of HSC-2 cells to *C. albicans* and *C. parapsilosis*

To examine the early effect of fungal presence on the metastatic cells, we aimed to characterize the host’s transcriptomic responses, that ultimately determines cellular functions. Thus, the transcriptome of HSC-2 cells was examined after *C. albicans* and *C. parapsilosis* stimuli at the early stages of the infection. Besides the comparative MOI of 1:1 applied in case of both species, we also selected a higher MOI for *C. parapsilosis* (MOI 5:1, pathogen:host), due to its less stimulatory nature.

After 6 hours of fungal presence significant host responses were triggered by both species, (FIG2), albeit most gene expression changes were due to *C. albicans*. Among the 170 differentially expressed genes (DEGs), 157 were significantly upregulated while 13 were downregulated (FIG2A, Table S1). Based GSEA results, the most relevant changes occurred within signaling pathways linked to inflammation. These included TNFa signaling via NFkB (NES: 3,82, padj: 2,51E-81), IL-2 -STAT5 signaling (NES: 2,79, padj: 1,98E-19), inflammatory signaling (NES: 2,95, padj: 4,56E-22) and hypoxia (NES: 2,99, padj: 9,80E-26) (FIG2B, Table S2). Besides inflammation, signaling pathways related to tumorous processes, e.g. epithelial-mesenchymal transition (NES: 2,82, padj: 6,89E-21), KRAS signaling (NES: 2,77, padj: 4,30E-17), MYC targets (NES: 2,69, padj: 8,61E-20) and p53 pathway (NES: 2,64, padj: 1,08E-17) were also markedly activated. Interestingly, these signaling pathways also exhibited a mild upregulation as early as 1 hour after the interaction (FIG S2B, Table S9). The top 20 DEGs with the highest expression variance among the treatment conditions are also linked to known inflammatory and cancerous functions (FIG2E, Table S3). Furthermore, all 20 have also been previously associated with oral cancer (Table S3). Under the applied conditions, the TOP20 DEGs were all significantly upregulated in the metastatic cell line during *C. albicans* exposure, while in case of *C. parapsilosis* presence their expression remained similar to that of the control condition.

**Figure 2.**
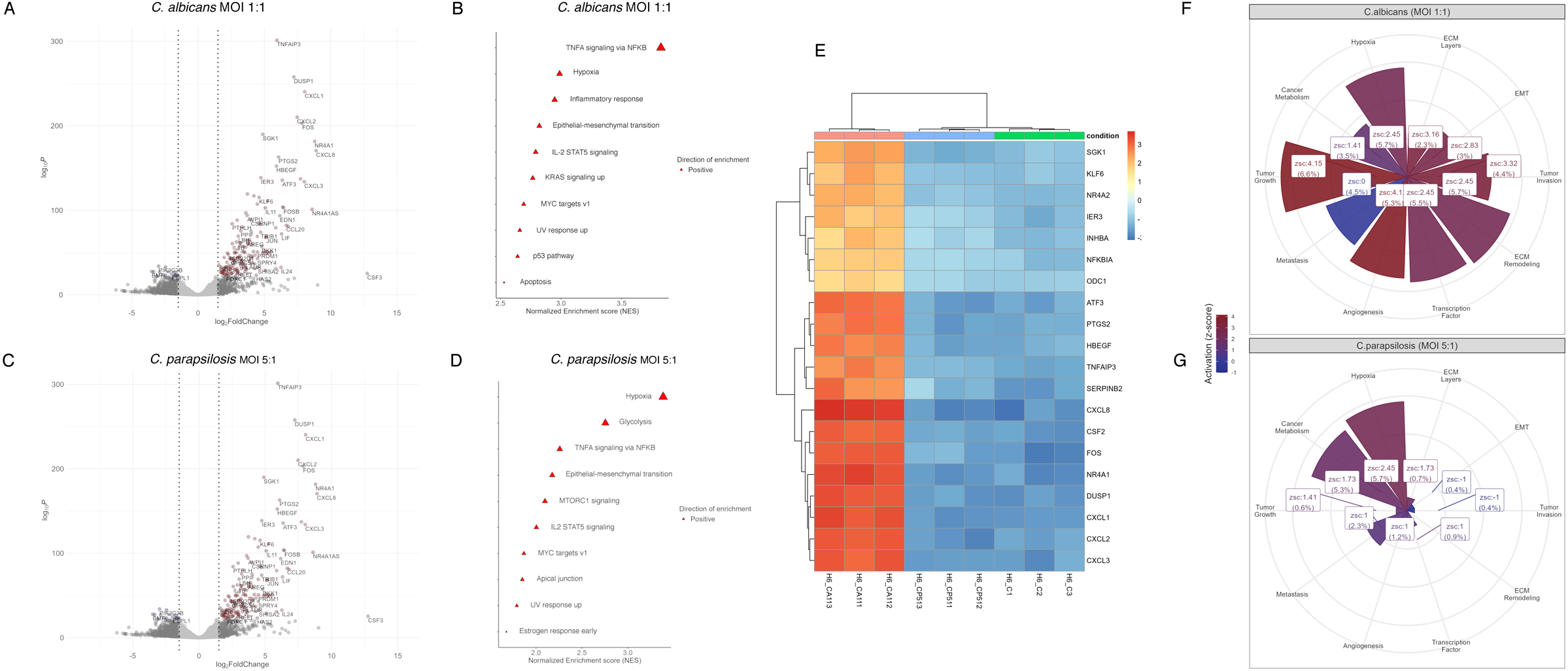
Early responses of HSC-2 cells to *C. albicans* and *C. parapsilosis* at the transcriptional level. Volcano plots represent significantly up-, and downregulated differentially expressed genes (DEGs) in HSC-2 cells, 6 hours after *C. albicans* (MOI 1:1, pathogen:host) (A), and high dose (C) *C. parapsilosis* (MOI 5:1, pathogen:host) stimuli. Gene-Set Enrichment Analyses (GSEA) highlight signaling pathways associated with the identified DEGs after exposure to *C. albicans* (B) and *C. parapsilosis* (D). The clustering heatmap (E) represents the top 20 DEGs with the highest expression variance among the treatment conditions. Charts to the right depict the tumor-associated categories of the PanCancer Progression Panel Gene list as represented by the identified DEGs after *C. albicans* (F) and *C. parapsilosis* (G) infection.

To further examine the potential pro-tumor effects of this species, we cross-referenced the identified DEGs with the nCounter® PanCancer Progression Panel Gene list provided by nanoString Technologies (openly accessible through nanoString). Our analyses revealed that 10 out of 10 tumor-associated categories are well represented by *C. albicans*-associated DEGs, and based on z-score values, 9 out of 10 groups exhibit widespread activation of the involved genes (FIG2F, Table S4). The most affected cancer hallmarks were tumor growth, ECM remodeling and hypoxia, based on the ratio of fungal-induced DEGs. These were followed by transcription factors, angiogenesis, metastasis, tumor invasion, cancer metabolism, EMT, and ECM layers. Supplementary figure S3 depicts how each of these categories are represented by the identified DEGs (FIG S3).

In contrast with *C. albicans*, *C. parapsilosis* triggered a less significant effect from HSC-2 cells, even at high fungal loads. Although differentially expressed genes were detected at the low infection dose - equivalent to that of *C. albicans -*, no significantly altered DEGs were identified. Contrarily, after the higher fungal load, 48 DEGs (46 upregulated, 2 downregulated) were identified (FIG2C, Table S5). GSEA analyses suggest their inclusion in pathways associated with inflammatory and tumor-associated pathways (TNFa signaling via NFKB (NES: 2,26, padj: 2,70E-09), hypoxia (NES: 3,39, padj: 3,89E-41), IL-2 STAT2 signaling (NES: 2,00, padj: 1,73E-05) and MYC targets v1 (NES: 1,87, padj: 4,01E-05), MTORC1 signaling (NES: 2,10, padj: 4,17E-08), epithelial-mesenchymal transition (NES: 2,18, padj: 1,62E-07), respectively), although based on normalized enrichment score values, these routes were less represented, than in case of *C. albicans* (FIG2D, Table S6). In cross-reference with the PanCancer Gene list, only the DEGs induced by the higher fungal load were linked to tumor-associated categories. Although these DEGs were found across 9 out of the 10 tumor groups, their representation was significantly lower compared to the *C. albicans* stimulus (FIG2G, Table S7).

Interestingly, according to our further analyses, a brief *C. albicans* stimulus, as short as 1 hour, was also able to induce, albeit a mild host response in HSC-2 cells. Under these conditions 4 DEGs were upregulated (FOS, EGR1, DUSP1 and CXCL8), all primarily linked to inflammation (FIG S2A-B, Tables S8-S9).

### Altered host miRome revealed during tumor-fungal interactions

Given the substantial transcriptional changes, next we examined how the developed responses are regulated. One potential mode of transcriptional regulation involves the action of microRNAs (miRs). As several miRs have already been described with a well-defined role in numerous biological pathways and molecular functions, we assessed the miR profile of HSC-2 cells after fungal stimuli.

Our results show that *C. albicans* significantly alters the miRome of the metastatic tumorous cell line, 6 hours after the co-incubation. We identified 23 differentially expressed miRs (DEmiRs), of which 8 showed up, while 15 showed downregulation (FIG3, Table S10). Interestingly, even a brief exposure to this species altered the expression of 3 miRs (miR-122-5p, miR-499a-5p and miR-152-3p, all upregulated) in HSC-2 cells (FIG S2C, Table S11).

**Figure 3.**
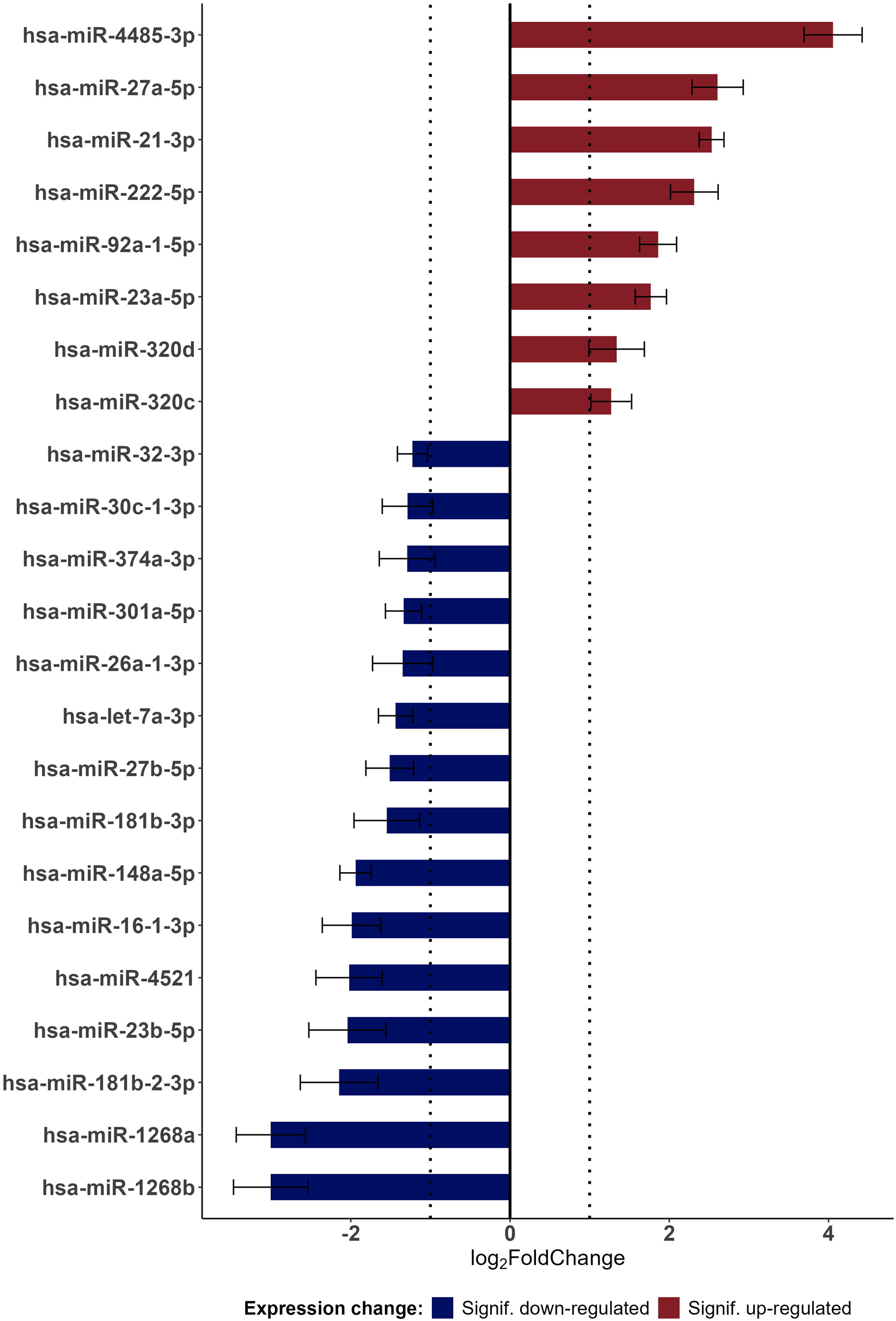
Altered miRome of HSC-2 cells after *C. albicans* stimulus. List of the identified DEmiRs in HSC-2 cells after 6 hours of *C. albicans* stimulus with the corresponding log2FoldChange values and adjusted p-values.

Contrarily to *C. albicans*, *C. parapsilosis* did not cause such a tremendous effect. Only one DEmiR, hsa-miR-210-5p (upregulated) was identified after the higher fungal load (FIG S4B, Table S12), while 6 showed altered expression (miR-3179 and miR-100-3p upregulated, miR-1-3p, miR-206, miR-6753-3p and miR-133a-3p all downregulated) following the lower fungal load (FIG S4A, Tables S13). Notably, all identified DEmiRs were specific to one condition only suggesting species-, dose- and time-dependency.

### *C. albicans*-induced miRs target inflammatory processes

To gain information about the potential function of DEmiRs, we aimed to identify their targets. Therefore, we cross-referenced the DEG lists with the corresponding DEmiR lists in each condition and predicted miR - target pairs.

In case of *C. parapsilosis* no potential miR-target pairs were identified, regardless of the fungal load. This suggests that although the expression of certain miRs is significantly altered in the presence of this species, their role in altering host regulatory mechanisms is negligible. Interestingly, the DEmiRs identified after brief exposure to *C. albicans* also lack predicted targets.

Contrarily, the DEmiRs identified after several hours of the *C. albicans* stimulus have numerous of targets. According to miR-target predictions, all the identified DEmiRs target more than 80% (143 of the total of 171) of the identified DEGs under the *C. albicans* condition, resulting in a total of 436 miR-target pairs (FIG4A, Table S14). This suggests the presence of a significant fungi-driven host response-regulatory effect. Using the recently updated miRTarBase (26), amongst these, we further identified 8 miR-target pairs (6 miRs targeting 8 genes) that had been experimentally validated (FIG4A, Table S14). These pairs were confirmed as functional miR-target interactions by previous studies. Of these, 1 pair (MAT2A - miR-21-3p) was validated by western-blot, reporter assays, qPCR and mircoarray - all suggesting a strong functional correlation -, while 7 pairs were validated via CLIP-Sequencing (PAR-CLIP or HITS-CLIP) (Table S14). Based on average expression levels at the miRNATissueAtlas2 (27) database, these miRs have also been detected from various surfaces, including squamous epithelial cells of mucous membranes (esophagus, tongue, oral cavity, saliva), as well as from tissues involved in immune responses (lymph nodes, salivary glands) (Table S14).

**Figure 4.**
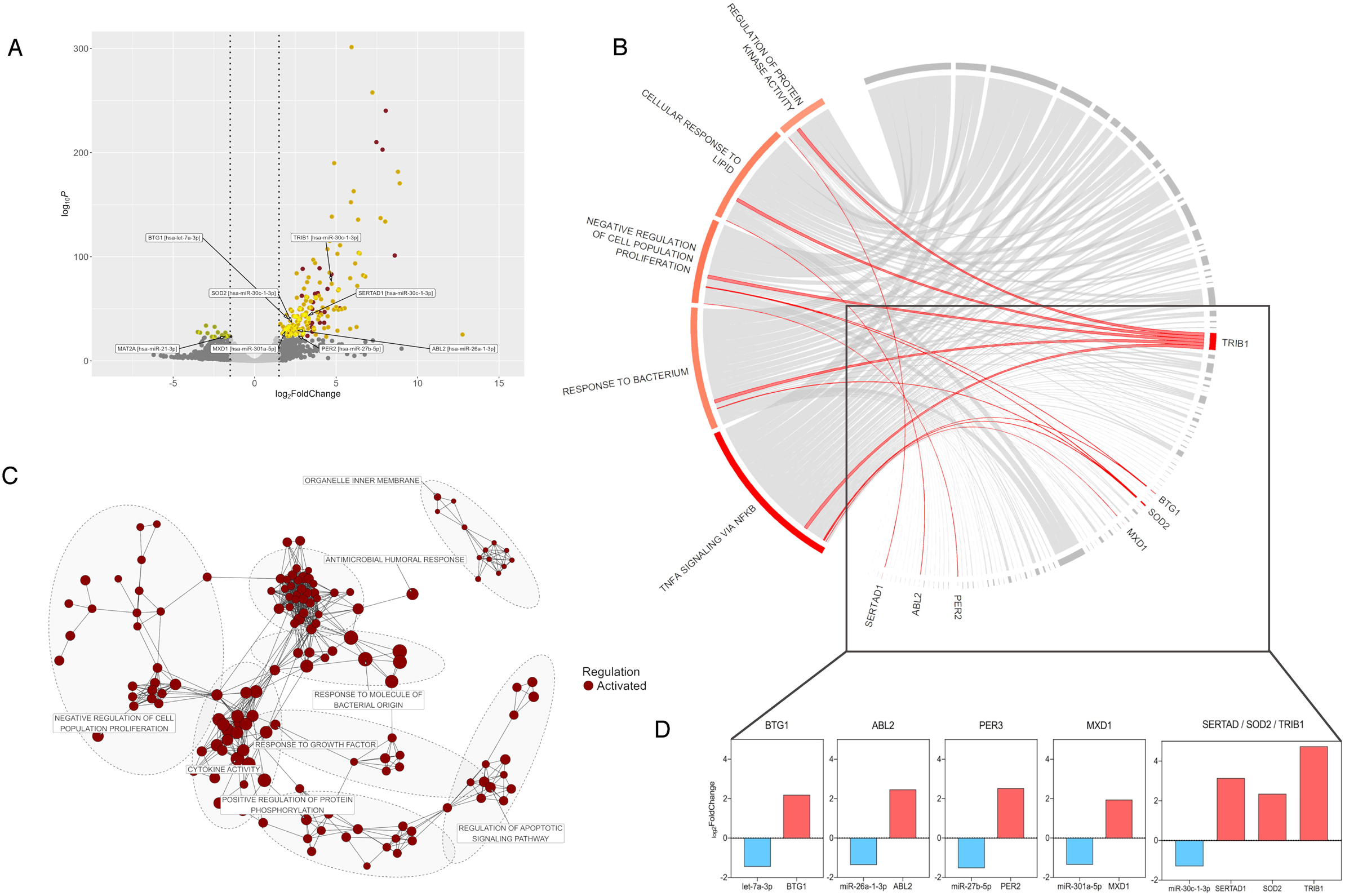
Potential functions of the *C. albicans*-induced miR-targets. (A) The volcano plot depicts the predicted targets of the 26 *C. albicans*-induced DEmiRs with yellow, and based on miRTarBase, the wet-lab validated targets with arrows and labels. (B) The network diagram shows the TOP200 GSEA clusters with potential functions associated with the miR targets. (C) Circo plot visualization of the TOP5 functions associated with the identified miR-target pairs with band width correspond to the calculated authority score of each target gene. (D) Highlighted, previously wet-lab validated miR-target pairs within the TOP5 functions. Expression values are derived from the results of the present study.

To gain an insight into the function of the miR targets, we first performed a network analysis based on all the corresponding GSEA results (FIG4B, Table S15). Subsequent results revealed ‘negative regulation of cell population proliferation’, cytokine activity’, ‘positive regulation of protein phosphorylation’, ‘response to growth factor’, ‘response to molecule of bacterial origin’, ‘regulation of apoptotic signaling pathway’, ‘antimicrobial humoral response’, and ‘organelle inner membrane’ as the top clusters where the identified DEGs accumulate after *C. albicans* infection. Next, we incorporated the predicted as well as the validated miR targets into the network analysis results, and sought to identify central functions, that are supported by the identified miR targets. As a result, ‘TNFA signaling via NFKB’ was identified as the primary function, supported by the miR targets, followed by pathways linked to ‘response to bacterium’, ‘negative regulation of cell proliferation’, ‘cellular responses to lipids’ and ‘regulation of protein kinase activity’ - suggesting that the fungi-driven host regulatory machinery also shifts host responses primarily towards pro-inflammation in the metastatic cell line (FIG4C, Table S16).

Seven functionally validated miR-target pairs are highlighted in these TOP functions (FIG4D). These include SERTAD1, SOD2, TRIB1 (all targeted by miR-30c-1-3p), ABL2 (miR-26a-1-3p), PER2 (miR-27b-5p), MXD1 (miR-301a-5p) and BTG1 (miR-let-7a-3p). While SERTAD1, ABL2, PER2 and MXD1 are each associated with only one category among the TOP5 categories, BTG1, SOD2 and TRIB1 are involved in multiple functions. Notably, 4 out of the 7 DEGs (MXD1, SOD2, BTG1 and TRIB1) are associated with ‘TNFA signaling via NFKB’, with TRIB1 being the most relevant, according to the centrality of their respective target genes in the network. Besides its highlighted role in NFKB signaling, TRIB1 is also strongly linked to the remaining TOP 4 pathways, suggesting a potentially central role for this gene. Interestingly, TRIB1 also has recognized oncogenic functions (28). Given TRIB1’s hereby recognized role in inflammation, and its known oncogenic functions, miR-30c-1-3p, a miRNA experimentally proven to target TRIB1, might potentially be used to silence both of its functions.

### *C. albicans*-influenced miRs further target tumor-associated genes

As the 7 highlighted miR targets have also been previously implicated in a number of malignancies (29–35) - albeit with controversial roles -, we aimed to further investigate how the altered miRome might affect tumor-associated functions. To achieve this, we cross-referenced the *C. albicans*-induced miRome list with the DEGs present in the nCounter® PanCancer Progression Panel Gene list.

As a result, we identified several miR-target pairs in each of the 10 cancer-associated categories (Table S17). By overlapping the miR-target pairs among the categories, we aimed to identify mutual DEGs that could warrant further attention. As a result, 23 DEGs were found with overlaps. Of these 2 were found to be present in 50%, 7 in 40%, and 6 in 30% of the cancer-associated categories (Table S18). The 2 DEGs with the highest overlap were JUN (L2FC: +5,16) and MMP3 (L2FC: +4,87), both of which have been previously suggested as markers of fungi-driven oral cancer progression (4) (FIG5A). These were followed by VEGFA (L2FC: +2,75), IL1B (L2FC: +2,59), SOX9 (L2FC: +2,79), EPHA2 (L2FC: +4,08), SDC4 (L2FC: +1,90), NR4A1 (L2FC: +8,79) and ADAMTS1 (L2FC: +3,94), several of which are also involved in inflammatory responses (36–42) (FIG5A). Their corresponding miRs are shown on FIG5B-C. The most represented categories by these genes are ‘Tumor growth’ and ‘Tumor invasion’, suggesting that these might be the primary mechanisms influenced by *C. albicans* during the early stages of fungi-driven OSCC progression. Among the identified miRs, miR-30c-1-3p targets 7 out of the 9 cancer-associated targets. Therefore, modulating the activity of this miR might also hold a therapeutic potential.

**Figure 5.**
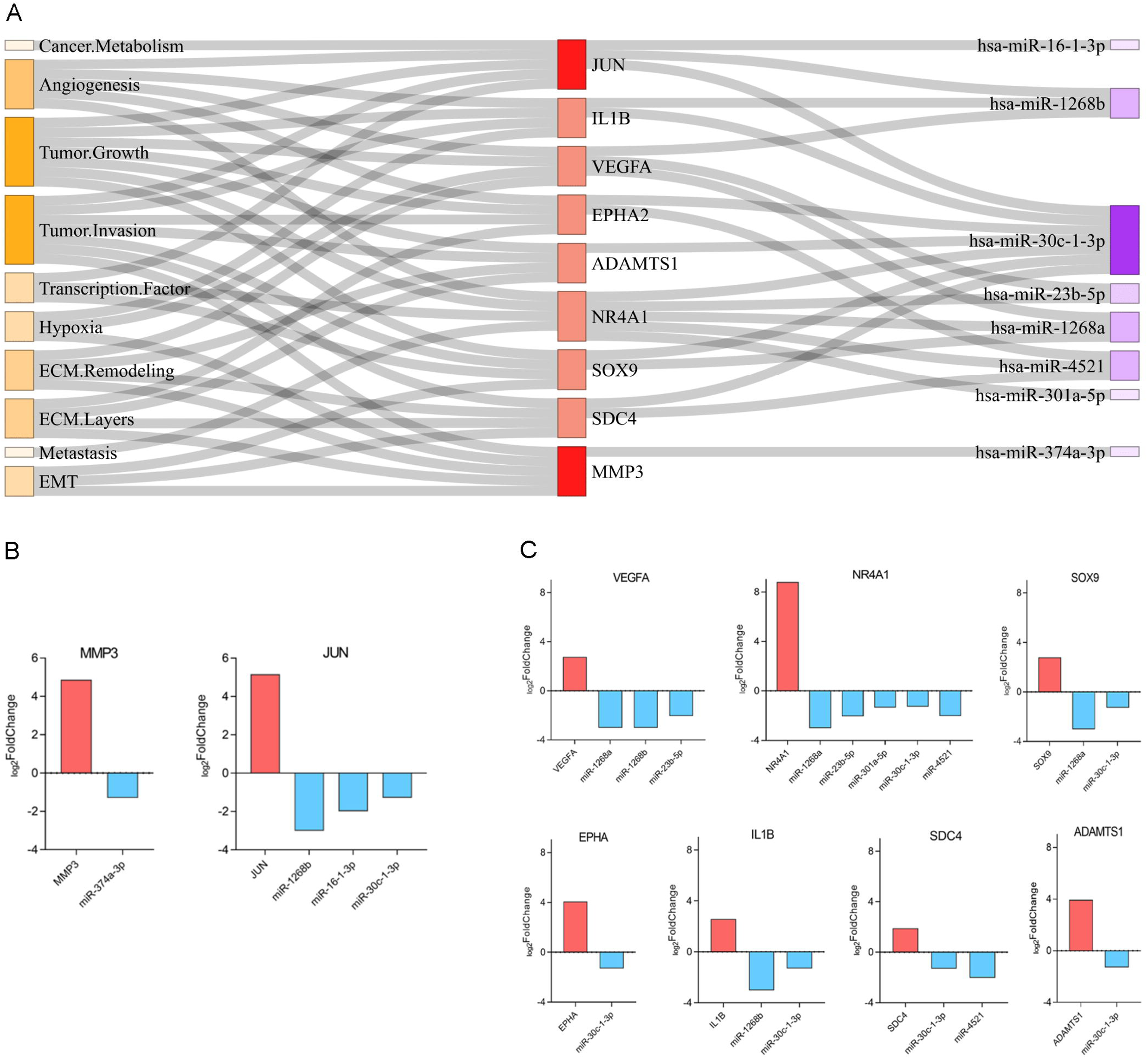
*C. albicans* - associated miR-targets with key roles in tumorigenic processes. (A) miR targets associated with tumorous processes, based on the PanCancer Progression Panel Gene list. The figure highlights miR-target genes, with the greatest overlap between the categories of the PanCancer Progression Panel and their correspondingly targeting miRs, bandwidths signifying regulation significance (negative log10(P-adj)), and facet colors regulation direction (L2FC, z-score). (B) JUN and MMP3, as genes present within 5 cancer-associated categories and their corresponding miRs. (C) Seven additional genes, present within 4 cancer-associated categories and their targeting miRs.

As the overexpression of both JUN and MMP3 has previously been implicated in *Candida*-driven tumor progression, their corresponding miRs (miR-1268b, miR-16-1-3p, miR-30c-1-3p against JUN and hsa-miR-374a-3p against MMP3) might serve as potentially valuable inhibitors.

## Discussion

Based on previous reports, the presence of oral candidiasis in patients with oral cancer is associated with high mortality (43). The association between oral squamous cell carcinoma (OSCC) and *Candida* presence has been recognized for nearly two decades, however, the causal relationship is still unclear. Recently, we demonstrated that the presence of *C. albicans* in fact acts as a direct driver of oral tumor progression, e.g. via enhancing MMP activity, the expression of tissue invasion markers, the production of oncometabolites and migration activity (4). Thus, we contextualized fungal-tumor interactions.

To deepen our understanding of fungi-driven oral tumor progression, we focused on examining the early stages of this interaction and by examining the simultaneously ongoing regulatory processes, we sought to identify key elements with potentially central functions, that might later possess translational potential.

Although several opportunistic pathogenic *Candida* species are present within the normal oral microbiota (44), *C. albicans* is the primarily species responsible for oral infections. While *C. parapsilosis* is the second most frequent *Candida* species within the normal oral flora, its presence is rarely associated with diseases. In our previous study conducted to compare the two species, we found that the two species triggered different host responses from healthy oral epithelial cells, that also reflected at the level of response regulation (miR level) (14). As even in the responses of normal oral epithelial cells we detected signaling pathways and post-transcriptional changes associated with tumorous processes, we proposed both species’ potential involvement in malignant transformations (14).

Our current findings indicate that even at the early stages of oral tumor-fungus interactions, both inflammatory responses and tumorous processes are detectable in the metastatic cell line, following exposure to either fungal species. However, these processes were significantly more pronounced in the presence of *C. albicans*.

Interestingly, a significant host response towards *C. parapsilosis* only occurred at the higher fungal load: hypoxia and glycolysis dominated host responses, similarly to what we have observed in normal oral epithelial cells (14). These were followed by the mild activation of pathways associated with inflammation (NFKB, IL2-STAT5 signaling) and tumor progression (e.g. EMT, MTORC, Myc targets). Since in our previous study, we did not reveal a role for this species in oral tumor progression *in vivo*, during prolonged colonization, the relevance of these findings during the early stages of the interaction remains to be elucidated. Interestingly, in healthy oral epithelial cells, this species significantly influenced processes related to vascular development and angiogenesis, regardless of the applied dose - an effect that was entirely absent in the current metastatic cell line. This suggests, that if *C. parapsilosis* has any relevant impact on tumorous processes, its effect might be stage-dependent and specific e.g. to the early stages of the disease. However, this idea needs to be further investigated.

During the early stages of tumor-fungal interactions, *C. albicans* induced mainly inflammatory responses from metastatic oral epithelial cells (TNFA signaling via NFKB), however, this was closely followed by the activation of tumor-associated pathways. In contrast, in normal oral epithelial cells, this species predominantly triggered almost exclusively inflammatory processes. Nevertheless, in two specific pathways—HIF1α signaling and hepatic fibrosis/stellate cell activation—*C. albicans* also clearly activated biological processes such as migration, angiogenesis, cell survival, and chemotaxis, suggesting its potential involvement in malignant transformation (14). The findings observed in both healthy and metastatic epithelial cell lines might indicate that *C. albicans* might play a role across multiple stages of the oral tumor progression. This hypothesis is supported by findings gained from a 4NQO animal model of carcinogenesis, where the presence of *C. albicans* enhanced dysplastic features in the tongue of mice (4).

In our study ‘KRAS_SIGNALING_UP’, ‘IL2_STAT5_SIGNALING’ and the ‘TNFA_SIGNALING_VIA_NFKB’ pathways were all upregulated by *C. albicans*. This is in contrast with a study by Hsieh et al. (8), where these pathways were predominantly upregulated in OSCC clinical specimens lacking *C. albicans* infection. This discrepancy might be attributed to differences in sample types and technical variables (e.g. length of fungal presence, amount of fungal burden, defined environmental conditions). Nonetheless, the highlight of these pathways in both studies suggests that they might represent key processes worthy of further investigations.

Interestingly, stratifin (SFN), identified as a potential biomarker of OSCC-coupled with *Candida* infection, did not appear in our list of DEGs under our experimental conditions. Nor did PD-L1, that has also been reported to be upregulated in other OSCC cell lines exposed to *C. albicans* (7). These discrepancies are likely due to the same factors discussed above. However, several other OSCC marker genes did show a significantly altered expression. These include IER3, INHBA, ATF3, HBEGF, FOS and CXCL1 - all of which were overexpressed.

Of these, EIR3 enhances the activation of wnt/β-catenin signaling (45), INHBA promotes proliferation, invasion and migration (46), ATF3 induces neoplastic lesions in the mouth of transgenic mice (47), while HBEGF contributes to invasion and MMP9 overexpression (48). Furthermore, CXCL1 overexpression is associated with poor prognosis, and also enhances invasion and migration of OSCC cells (49), while FOS is suggested to play a role in cancer stem cell reprogramming in HNSCC (50). Interestingly, other OSCC-associated biomarkers, albeit with a suppressor role, were also upregulated after *C. albicans* presence. These include KLF6 (51), SERPINB2 (52), NR4A1 (53), or DUSP1 (54). Although the expression of these markers is highly influenced by experimental or environmental conditions, further investigations are needed to answer these discrepancies.

Alterations in miR function have been associated with various diseases, including cancer (12). Several miRs have also already been implicated in oral cancer (55). We hence investigated how the presence of fungal cells influences the miR response of the metastatic cell line, and how these alterations may contribute to fungi-driven OSCC progression. We identified several differentially expressed miRs across the different treatment conditions. Notably, there was no overlap in miRs between the conditions, indicating that their expression in species-, dose- and time-dependent - similarly to what we observed in normal oral epithelial cell responses (14). Furthermore, when comparing the miR profiles of the metastatic cell line to those of healthy oral epithelial cells, we found only minimal overlap. This suggests that the physiological status of the mucosal epithelium also influences the miR response to fungi. Moreover, the expression of the 4 overlapping miRs (miR-92a, miR-16, hsa-miR-181b and hsa-miR-4485-3p) changed in opposite directions between the two host conditions when exposed to *C. albicans* (14).

The majority of miR expression changes were induced by *C. albicans*. Interestingly, although no significant transcriptional changes were detected after the low dose *C. parapsilosis* stimulus, 6 miRs still showed significantly altered expression. Among these, 4 miRs have been previously associated with OSCC (56–59). However, as no corresponding DEGs could be linked to these miRs, the relevance of this effect is likely negligible. Similarly, the higher fungal load affected the expression of only 1 miR, albeit with no corresponding DEGs.

Of the 23 miRs identified after *C. albicans* infection, 17 have previously been linked to OSCC. Their expression in OSCC is also well-characterized. In the presence of the fungus, a smaller subset of these miRs exhibited changes in the opposite direction. Specifically, *C. albicans* induced opposite directional changes in the expression of 5 OSCC-linked miRs (miR-181b, miR-301a, miR-32, miR-320, miR-4485-3p) (60–63). The significance of this phenomenon remains to be elucidated. Contrarily, for a significant proportion of the identified OSCC-miRs (12 miRs), the fungus caused an amplification effect, and promoted their expression in the same direction. Hsa-let-7a, miR-148a, miR-16, miR-26, miR-374a, miR-23b and miR-27b are suggested tumor suppressors in OSCC. Their expression in oral cancer is downregulated (64–70). Here, *C. albicans* further reduced their expression. Although their role in OSCC is not yet clarified, miR-4521 and miR-30c-1-3-p - both known tumor suppressors in pancreatic, lung and breast cancer (71–73) - also showed reduced expression in response to *C. albicans*. miR-21-3p, miR-222, miR-23a, miR-27a and miR-92a are all upregulated, hence oncogenic miRs in oral cancer (74–78). In this study, *C. albicans* further increased their expression. Thus, *C. albicans* affected the expression of several OSCC-associated miRs, by further inhibiting tumor suppressor miRs, while promoting oncogenic miRs.

Subsequent miR-target analyses revealed, that the identified miRs regulate more than 80% of the *C. albicans*-induced DEGs. These miR targets are primarily linked to inflammatory processes. This finding is consistent with the literature, suggesting that tumor-microbe interactions are predominantly mediated through inflammatory pathways (79–81). Based on our predictions, the key miR targets linking the main functional pathways are SERTAD1, PER2, ABL2, MXD1, SOD2, BTG1 and TRIB1. As interestingly, all 7 genes have previously been implicated in a number of malignancies - some also in HNSCC - (29–35), and their corresponding miRs are also strongly associated with OSCC regulation (62, 64, 66, 68, 71), these miR-target pairs may be valuable candidates for future investigations. Our predictions suggest that TRIB1 is also associated with all TOP 5 functional categories, indicating that it may be a potentially central gene in *C. albicans*-driven regulatory processes. It has also been suggested that TRIB1 may inactivate p53 function and activate oncogenic Pi3K-AKT and MAPK signaling pathways (28). *C. albicans* is also well-known to interfere with Pi3K-AKT and MAPK signaling in mucosal cells (82, 83), which could provide a commons basis for mechanisms ultimately leading to progression events. Therefore, further investigating the TRIB1 - miR-30c-1-3p connection might hold a valuable potential.

Next we investigated the miR-mediated regulation of PanCancer Progression genes. We identified several PanCancer genes with an altered expression in the metastatic cells, in response to *C. albicans*. Notably, many of these genes were also found to be targets of the *C. albicans*-induced miRs. Among these, 29 genes overlapped in at least 2 of the 10 designated categories. The greatest overlap between the categories was observed for JUN and MMP3 - with both being present in 4 categories. JUN is a component of the *C. albicans*-induced mucosal epithelial immune response, and its transient activation is primarily observed after the adhesion of the yeast form to epithelial cells, mediated through JNK/ERK1/2 signaling (84). Although its activation has yet unknown effects (85), we have previously suggested its involvement in *C. albicans*-driven OSCC progression due to its marked overexpression (4). Besides JUN, we further demonstrated the significant overactivation of MMP3 - along with other matrix metalloproteases - in metastatic oral epithelial cells when the fungus is present on the long term (4). As both genes are already implicated in the *C. albicans*-driven oral tumor progression, their targeting with their corresponding miRs might also hold a valuable potential. Among the remaining 7 PanCancer progression markers with a high overlap, the role of 3 (EPHA2, IL1B and VEGFA) has also been addressed in the context of *C. albicans*-oral epithelial cell interactions. The accumulation of Epha2 - an epithelial pattern recognition receptor - was observed in the presence of *C. albicans* (86), while IL1B - a proinflammatory cytokine - was shown to be overexpressed as part of an antifungal response against this species (87). VEGFA is also overexpressed in metastatic oral epithelial cells as well as in melanoma cells exposed to this species (4, 88).

The expression of NR4A1 and SCD4 has not yet been addressed in epithelial cells in the context of *C. albicans* exposure. Recent studies indicate downregulation of NR4A1 in the colon of *C. albicans*-colonized mice (89). A potential role has also been suggested for SDC4 during the adhesion of *C. albicans* to the corneal epithelium (90). ADAMTS1 and SOX9 expression during *C. albicans* presence has not yet been investigated in any model. Each of the hereby identified genes potentially linked to fungi-driven OSCC progression might also be targeted through their corresponding inhibitory miRs identified in this study.

To summarize, in this study, we aimed to investigate the fungi-driven tumor progression phenomenon in more depth. We focused on exploring the regulatory processes behind this phenomenon, at the early stages of the tumor-fungal interactions. Through this approach, we not only added an additional layer to the understanding of this event, but also identified regulatory elements that might offer new perspectives for potential interventions. Our analyses highlighted several miR-target pairs, among which TRIB - miR-30c-1-3p, JUN - miR-16-1-3p/miR-1268b/miR-30c-1-3p and MMP3 - miR-374a-3p appear most promising. Thus, our findings demonstrate, that the previously observed ability of *C. albicans* to promote oral tumor progression is also evident at the miR level. In this context, the fungus might further enhance pro-tumor processes by altering post-transcriptional regulatory mechanisms.

## Materials and Methods

### Applied strains and their cultivation

This study utilized *Candida albicans* SC5314 and *Candida parapsilosis* CLIB 214 strains. The fungal strains were maintained on solid YPD medium supplemented with 1% penicillin-streptomycin (PS) at 4°C. Yeast cultures were grown overnight in liquid YPD medium at 30°C, washed three times with phosphate-buffered saline (1x PBS) and adjusted to the desired concentration.

### Epithelial cells and their stimulation

Cells of the OSCC cell line, HSC-2, were cultured in Eagle’s minimum essential medium (EMEM) supplemented with 10% heat inactivated FBS, 1% PS, and 4 mM L-glutamine. Cells were handled and maintained according to the manufacturer’s suggestions at 37°C in the presence of 5% CO_2_. Cells were then stimulated with fungi at the desired multiplicity of infection ratios (MOIs as pathogen: host), in serum-free media, at 37°C in the presence of 5% CO_2_. Following stimuli, either host cells or cell-free supernatants were collected for the subsequent experiment. For qPCR analysis, host cells were stimulated with fungal cells for 1 hour or 6 hours, while for ELISA assays for 24 hours.

### Extraction of total RNA and miRNA

Total RNAs and miRNAs were extracted from epithelial cells using the miRNeasy Mini Kit, following the manufacturer’s instructions with slight modifications. Epithelial cells were cultured in supplemented EMEM medium in six-well tissue culture plates until they reached 85% confluency, then washed once with PBS and stimulated with *C. albicans* (MOI 1:1) or *C. parapsilosis* (MOI 1:1 and/or 5:1) in unsupplemented EMEM/RPMI medium. After coincubation, the culture medium and any floating fungal cells were removed, and host cells were washed twice with 1xPBS before lysis with the QIAzol lysis reagent. Plates were gently agitated. Following lysis, phase separation, and RNA purification, the samples underwent quality and quantity assessments before cDNA synthesis or library preparation.

### cDNA synthesis and real-time PCR

1000Lng of RNA was used for cDNA synthesis using the RevertAid first strand cDNA synthesis kit. cDNS samples were prepared according to the manufacturer’s guidelines. For qPCR analyses the following real time primers were used: 5’-GGAGCGCCTGATAATCCAGT-3’ forward (fw) and 5’-ATCTGTCACGTTCTTGGGGC-3’ reverse (rv) for JUN, 5’-AACACATTAGACGGCCTCCTG-3’ fw and 5’-CTGTTGATCTGGGTCTTGGCT-3’ rv for LAMC2, 5’-CAGAGATGAAGTCCGGTTTTTC-3’ fw and 5’-GGGGTATCCGTGTAGCACAT-3’ rv for MMP1, 5’-GCTCTGCCTATCCTCTGAGTG-3’ fw and 5’-CAACGTCAGGAACTCCACACC-3’ rv for MMP10, and 5’-GTGCCAATACCATGAAGAGGA-3’ fw and 5’-CTCTTTCTGGTCCCCACTCTT-3’ rv for INHBA. The reaction was carried out in a C1000 thermal cycler under the following conditions: 95°C for 3 minutes, followed by 95°C for 10s, then 50 cycles with 60°C for 30 seconds, and 65°C for 30 seconds, finally by 72°C for 30s. The fold change in mRNA expression was determined using the ΔΔCT method (threshold cycle), with β2-Microglobulin serving as internal housekeeping gene control.

### Cytokine Assays

The concentration of cytokines were measured with commercial enzyme-linked immunosorbent assay (ELISA) kits after the 24-hour fungal infection of epithelial cells at 37°C in the presence of 5% CO2. All assays were performed according to the manufacturers’ instructions.

### miRNA/RNA sequencing library preparation and sequencing

m(i)RNA sequencing libraries were prepared at Novogene (UK) Company Limited, Cambridge, UK. For RNA sequencing mRNA was enriched using poly(A) capture during library preparation, and sequencing was performed on an Illumina NovaSeq X Plus platform with a 2 × 150 bp paired-end configuration, following the NovaSeq PE150 workflow. The target sequencing depth aimed to achieve an average of ≥ 20 million read pairs per sample. For miRNA sequencing small RNA libraries were prepared and sequencing was performed on an Illumina NovaSeq 6000 platform with single-end 50 bp configuration, following the NovaSeq SE50 workflow. Target sequencing depth aimed to achieve an average of ≥ 10 million read pairs per sample.

### Transcriptome analysis

Preliminary quality analysis and trimming of raw sequence files were performed using trim galore (v0.6.4) with quality Phred score cutoff 20, and Illumina adapter for trimming. Reads were then aligned to the GRCh38 reference genome with HISAT2(v2.2.1), with parameters -- dta --non-deterministic --rna-strandness, aimed for consequential quantification using FeatureCounts(v2.0.3), performed at the gene level. Next, differential gene expression analysis (logarithmic fold change, LFC) was conducted using DESeq2(v1.46.0). Low-expression genes (<1 ppm in at least 3 samples) were filtered out, and DEGs were identified based on an absolute logaritmic2 fold change (L2FC) >1.5 and an adjusted P-adj value <0.05. Multiple testing correction was performed via the fdrtools package(v.1.2.18) applying the Benjamini and Hochberg (BH) method. Principal component analysis (PCA) was conducted to assess sample clustering, and volcano plots and heatmaps of significant genes were generated to visualize expression patterns via the ggplot2 (v3.5.1) and pheatmap (v1.0.12) packages, respectively.

### miRNA mapping and count

Sequenced reads were mapped to known and novel microRNA precursors from miRBase (version 22) using miRDeep2.0 (v2.0.1.3). Hits with read counts below 1 ppm in at least three samples were excluded from further analysis. To differential expression was computed by the DESeq2 package, and P-values were adjusted via the fdrtools package, correspondingly to the transcriptome analysis. Significant hits were defined by a P-adj <0.05 and an absolute L2FC > 1.0. Possible miRNA interactions were identified through target mining the miRWalk2 database (91). Interaction sites across all regions of a gene were considered(score > 0.95), while an available miRTarBase(release 9) accession was required for validation.

### Pathway analyses

Following genome-wide RNA and miRNA expression analyses, gene expression data were interpreted using gene set enrichment analysis(GSEA) via the fgsea(v1.32.2). Both analyses were carried out against a constant background, using the human genome-wide annotation package org.Hs.eg.db(v3.20.0), filtered for the genes actively expressed in the OSCC samples. An activation z-score was integrated with the enrichment results, and for further data mining network analysis was conducted. Briefly, an adjacency matrix was created including all GO terms and KEGG pathways, based on Cohen’s kappa similarities with cutoff>0.25. The network was then generated from the adjacency matrix, and clusters were detected using a greedy, Louvain community detection algorithm. For determining cluster representative terms the centrality of the nodes was calculated, weighted on gene ration and expression pattern of the involved genes, and terms with the highest hub score were selected as cluster representatives.

### Statistical analysis

Statistical analyses were conducted using GraphPad Prism v6.0 software, applying parametric t-tests. Differences were considered statistically significant at P < 0.05.

## Data availability

Sequencing data are accessible under the BioProject accession number GSE296481.

## Author contributions

R.T. and A.G. contributed to the concept and design of the project. R.T. carried out the experiments. M.H. analyzed the acquired data and generated the figures with the help of R.T and J.N. R.T. prepared the original manuscript and A.G., J.N, and M.H. reviewed, discussed, and approved the final version of the manuscript.

## References

1. Kandalai S, Li H, Zhang N, Peng H, Zheng Q. 2023. The human microbiome and cancer: a diagnostic and therapeutic perspective. Cancer Biol Ther 24:2240084.

2. Newton R, de Martel C, Plummer M. 2022. Infectious agents: Missed opportunities for prevention.World Cancer Report: Cancer research for cancer prevention. Lyon (FR).

3. Hosseini K, Ahangari H, Chapeland-Leclerc F, Ruprich-Robert G, Tarhriz V, Dilmaghani A. 2022. Role of Fungal Infections in Carcinogenesis and Cancer Development: A Literature Review. Adv Pharm Bull 12:747–756.

4. Vadovics M, Ho J, Igaz N, Alföldi R, Rakk D, Veres É, Szücs B, Horváth M, Tóth R, Szücs A, Csibi A, Horváth P, Tiszlavicz L, Vágvölgyi C, Nosanchuk JD, Szekeres A, Kiricsi M, Henley-Smith R, Moyes DL, Thavaraj S, Brown R, Puskás LG, Naglik JR, Gácser A. 2021. Candida albicans Enhances the Progression of Oral Squamous Cell Carcinoma In Vitro and In Vivo. mBio 13:e0314421.

5. İlhan B, Vural C, Gürhan C, Vural C, Veral A, Wilder-Smith P, Özdemir G, Güneri P. 2023. Real-Time PCR Detection of Candida Species in Biopsy Samples from Non-Smokers with Oral Dysplasia and Oral Squamous Cell Cancer: A Retrospective Archive Study. Cancers (Basel) 15:5251.

6. Lee Y-H, Jung J, Hong J-Y. 2024. Oral Microbial Changes in Oral Squamous Cell Carcinoma: Focus on Treponema denticola, Lactobacillus casei, and Candida albicans. Medicina (Kaunas) 60:1753.

7. Wang X, Zhao W, Zhang W, Wu S, Yan Z. 2022. Candida albicans induces upregulation of programmed death ligand 1 in oral squamous cell carcinoma. J Oral Pathol Med 51:444–453.

8. Hsieh Y-P, Wu Y-H, Cheng S-M, Lin F-K, Hwang D-Y, Jiang S-S, Chen K-C, Chen M-Y, Chiang W-F, Liu K-J, Huynh NC-N, Huang W-T, Huang T-T. 2022. Single-Cell RNA Sequencing Analysis for Oncogenic Mechanisms Underlying Oral Squamous Cell Carcinoma Carcinogenesis with Candida albicans Infection. Int J Mol Sci 23:4833.

9. Di Leva G, Garofalo M, Croce CM. 2014. MicroRNAs in cancer. Annu Rev Pathol 9:287–314.

10. Saliminejad K, Khorram Khorshid HR, Soleymani Fard S, Ghaffari SH. 2019. An overview of microRNAs: Biology, functions, therapeutics, and analysis methods. J Cell Physiol 234:5451–5465.

11. Ho PTB, Clark IM, Le LTT. 2022. MicroRNA-Based Diagnosis and Therapy. Int J Mol Sci 23:7167.

12. Peng Y, Croce CM. 2016. The role of MicroRNAs in human cancer. Signal Transduct Target Ther 1:15004.

13. 2024. What will it take to get miRNA therapies to market? Nat Biotechnol 42:1623– 1624.

14. Horváth M, Nagy G, Zsindely N, Bodai L, Horváth P, Vágvölgyi C, Nosanchuk JD, Tóth R, Gácser A. 2021. Oral Epithelial Cells Distinguish between Candida Species with High or Low Pathogenic Potential through MicroRNA Regulation. mSystems 6:e00163–21.

15. Tasso CO, Ferrisse TM, de Oliveira AB, Ribas BR, Jorge JH. 2023. Candida species as potential risk factors for oral squamous cell carcinoma: Systematic review and meta-analysis. Cancer Epidemiol 86:102451.

16. Ayuningtyas NF, Mahdani FY, Pasaribu TAS, Chalim M, Ayna VKP, Santosh ABR, Santacroce L, Surboyo MDC. 2022. Role of Candida albicans in Oral Carcinogenesis. Pathophysiology 29:650–662.

17. Mäkinen AI, Mäkitie A, Meurman JH. 2021. Candida prevalence in saliva before and after oral cancer treatment. Surgeon 19:e446–e451.

18. Abidullah M, Bhosle S, Komire B, Sharma P, Swathi K, Karthik L. 2021. Investigation of Candidal Species among People Who Suffer from Oral Potentially Malignant Disorders and Oral Squamous Cell Carcinoma. J Pharm Bioallied Sci 13:S1050–S1054.

19. Sankari SL, Mahalakshmi K, Kumar VN. 2020. A comparative study of Candida species diversity among patients with oral squamous cell carcinoma and oral potentially malignant disorders. BMC Res Notes 13:488.

20. Roy SK, Astekar M, Sapra G, Chitlangia RK, Raj N. 2019. Evaluation of candidal species among individuals with oral potentially malignant disorders and oral squamous cell carcinoma. J Oral Maxillofac Pathol 23:302.

21. Hulimane S, Maluvadi-Krishnappa R, Mulki S, Rai H, Dayakar A, Kabbinahalli M. 2018. Speciation of Candida using CHROMagar in cases with oral epithelial dysplasia and squamous cell carcinoma. J Clin Exp Dent 10:e657–e660.

22. Perera M, Al-Hebshi NN, Perera I, Ipe D, Ulett GC, Speicher DJ, Chen T, Johnson NW. 2017. A dysbiotic mycobiome dominated by Candida albicans is identified within oral squamous-cell carcinomas. J Oral Microbiol 9:1385369.

23. Sanketh DS, Patil S, Rao RS. 2016. Estimating the frequency of Candida in oral squamous cell carcinoma using Calcofluor White fluorescent stain. J Investig Clin Dent 7:304–307.

24. Jahanshahi G, Shirani S. 2015. Detection of Candida albicans in oral squamous cell carcinoma by fluorescence staining technique. Dent Res J (Isfahan) 12:115–120.

25. Berkovits C, Tóth A, Szenzenstein J, Deák T, Urbán E, Gácser A, Nagy K. 2016. Analysis of oral yeast microflora in patients with oral squamous cell carcinoma. Springerplus 5:1257.

26. Cui S, Yu S, Huang H-Y, Lin Y-C-D, Huang Y, Zhang B, Xiao J, Zuo H, Wang J, Li Z, Li G, Ma J, Chen B, Zhang H, Fu J, Wang L, Huang H-D. 2025. miRTarBase 2025: updates to the collection of experimentally validated microRNA-target interactions. Nucleic Acids Res 53:D147–D156.

27. Keller A, Gröger L, Tschernig T, Solomon J, Laham O, Schaum N, Wagner V, Kern F, Schmartz GP, Li Y, Borcherding A, Meier C, Wyss-Coray T, Meese E, Fehlmann T, Ludwig N. 2022. miRNATissueAtlas2: an update to the human miRNA tissue atlas. Nucleic Acids Res 50:D211–D221.

28. Singh K, Showalter CA, Manring HR, Haque SJ, Chakravarti A. 2024. “Oh, Dear We Are in Tribble”: An Overview of the Oncogenic Functions of Tribbles 1. Cancers (Basel) 16:1889.

29. Nguyen HA, Vu SH, Jung S, Lee BS, Nguyen TNQ, Lee H, Lee H-G, Myagmarjav D, Jo T, Choi Y, Lee M-S. 2022. SERTAD1 Sensitizes Breast Cancer Cells to Doxorubicin and Promotes Lysosomal Protein Biosynthesis. Biomedicines 10:1148.

30. Jones JK, Zhang H, Lyne A-M, Cavalli FMG, Hassen WE, Stevenson K, Kornahrens R, Yang Y, Li S, Dell S, Reitman ZJ, Herndon JE, Hoj J, Pendergast AM, Thompson EM. 2023. ABL1 and ABL2 promote medulloblastoma leptomeningeal dissemination. Neurooncol Adv 5:vdad095.

31. Ao Y, Zhao Q, Yang K, Zheng G, Lv X, Su X. 2018. A role for the clock period circadian regulator 2 gene in regulating the clock gene network in human oral squamous cell carcinoma cells. Oncol Lett 15:4185–4192.

32. Du F, Dong D, Zhang X, Jia J. 2021. MXD1 is a Potential Prognostic Biomarker and Correlated With Specific Molecular Change and Tumor Microenvironment Feature in Esophageal Squamous Cell Carcinoma. Technol Cancer Res Treat 20:15330338211052142.

33. Ding D, Li N, Ge Y, Wu H, Yu J, Qiu W, Fang F. 2024. Current status of superoxide dismutase 2 on oral disease progression by supervision of ROS. Biomed Pharmacother 175:116605.

34. Li Y, Huo J, He J, Zhang Y, Ma X. 2020. BTG1 inhibits malignancy as a novel prognosis signature in endometrial carcinoma. Cancer Cell Int 20:490.

35. Kim T, Johnston J, Castillo-Lluva S, Cimas FJ, Hamby S, Cardiogenics Consortium*, Gonzalez-Moreno S, Villarejo-Campos P, Goodall AH, Velasco G, Ocana A, Muthana M, Kiss-Toth E. 2022. TRIB1 regulates tumor growth via controlling tumor-associated macrophage phenotypes and is associated with breast cancer survival and treatment response. Theranostics 12:3584–3600.

36. Lai WK, Adams DH. 2005. Angiogenesis and chronic inflammation; the potential for novel therapeutic approaches in chronic liver disease. J Hepatol 42:7–11.

37. Ren K, Torres R. 2009. Role of interleukin-1beta during pain and inflammation. Brain Res Rev 60:57–64.

38. Ouyang Y, Wang W, Tu B, Zhu Y, Fan C, Li Y. 2019. Overexpression of SOX9 alleviates the progression of human osteoarthritis in vitro and in vivo. Drug Des Devel Ther 13:2833–2842.

39. Finney AC, Funk SD, Green JM, Yurdagul A, Rana MA, Pistorius R, Henry M, Yurochko A, Pattillo CB, Traylor JG, Chen J, Woolard MD, Kevil CG, Orr AW. 2017. EphA2 Expression Regulates Inflammation and Fibroproliferative Remodeling in Atherosclerosis. Circulation 136:566–582.

40. Giuliani A, Ramini D, Sbriscia M, Crocco P, Tiano L, Rippo MR, Bonfigli AR, Rose G, De Luca M, Olivieri F, Sabbatinelli J. 2024. Syndecan 4 is a marker of endothelial inflammation in pathological aging and predicts long-term cardiovascular outcomes in type 2 diabetes. Diabetol Metab Syndr 16:203.

41. McMorrow JP, Murphy EP. 2011. Inflammation: a role for NR4A orphan nuclear receptors? Biochem Soc Trans 39:688–693.

42. Krampert M, Kuenzle S, Thai SN-M, Lee N, Iruela-Arispe ML, Werner S. 2005. ADAMTS1 proteinase is up-regulated in wounded skin and regulates migration of fibroblasts and endothelial cells. J Biol Chem 280:23844–23852.

43. Di Cosola M, Cazzolla AP, Charitos IA, Ballini A, Inchingolo F, Santacroce L. 2021. Candida albicans and Oral Carcinogenesis. A Brief Review. J Fungi (Basel) 7:476.

44. Ghannoum MA, Jurevic RJ, Mukherjee PK, Cui F, Sikaroodi M, Naqvi A, Gillevet PM. 2010. Characterization of the oral fungal microbiome (mycobiome) in healthy individuals. PLoS Pathog 6:e1000713.

45. Yin C, Miao Y, Lu W, Liu Z. 2025. IER3 Facilitates Tumor Progression and Aberrant Glycolysis via Activating wnt/β-Catenin Pathway in Oral Squamous Cell Carcinoma. Adv Biol (Weinh) e2400564.

46. Chen X, Fan R. 2024. Inhibin A contributes to the tumorigenesis of oral squamous cell carcinoma by KIAA1429-mediated m6A modification. J Oral Pathol Med 53:266–274.

47. Wang A, Arantes S, Conti C, McArthur M, Aldaz CM, MacLeod MC. 2007. Epidermal hyperplasia and oral carcinoma in mice overexpressing the transcription factor ATF3 in basal epithelial cells. Mol Carcinog 46:476–487.

48. Ohnishi Y, Inoue H, Furukawa M, Kakudo K, Nozaki M. 2012. Heparin-binding epidermal growth factor-like growth factor is a potent regulator of invasion activity in oral squamous cell carcinoma. Oncol Rep 27:954–958.

49. Wei L-Y, Lee J-J, Yeh C-Y, Yang C-J, Kok S-H, Ko J-Y, Tsai F-C, Chia J-S. 2019. Reciprocal activation of cancer-associated fibroblasts and oral squamous carcinoma cells through CXCL1. Oral Oncol 88:115–123.

50. Muhammad N, Bhattacharya S, Steele R, Phillips N, Ray RB. 2017. Involvement of c-Fos in the Promotion of Cancer Stem-like Cell Properties in Head and Neck Squamous Cell Carcinoma. Clin Cancer Res 23:3120–3128.

51. Hsu L-S, Huang R-H, Lai H-W, Hsu H-T, Sung W-W, Hsieh M-J, Wu C-Y, Lin Y-M, Chen M-K, Lo Y-S, Chen C-J. 2017. KLF6 inhibited oral cancer migration and invasion via downregulation of mesenchymal markers and inhibition of MMP-9 activities. Int J Med Sci 14:530–535.

52. Huang Z, Li H, Huang Q, Chen D, Han J, Wang L, Pan C, Chen W, House MG, Nephew KP, Guo Z. 2014. SERPINB2 down-regulation contributes to chemoresistance in head and neck cancer. Mol Carcinog 53:777–786.

53. Wang Y, Ren X, Li W, Cao R, Liu S, Jiang L, Cheng B, Xia J. 2021. SPDEF suppresses head and neck squamous cell carcinoma progression by transcriptionally activating NR4A1. Int J Oral Sci 13:33.

54. Zhang X, Hyer JM, Yu H, D’Silva NJ, Kirkwood KL. 2014. DUSP1 phosphatase regulates the proinflammatory milieu in head and neck squamous cell carcinoma. Cancer Res 74:7191–7197.

55. Wang J, Lv N, Lu X, Yuan R, Chen Z, Yu J. 2021. Diagnostic and therapeutic role of microRNAs in oral cancer (Review). Oncol Rep 45:58–64.

56. He B, Lin X, Tian F, Yu W, Qiao B. 2018. MiR-133a-3p Inhibits Oral Squamous Cell Carcinoma (OSCC) Proliferation and Invasion by Suppressing COL1A1. J Cell Biochem 119:338–346.

57. Wang L, Song H, Yang S. 2021. MicroRNA-206 has a bright application prospect in the diagnosis of cases with oral cancer. J Cell Mol Med 25:8169–8173.

58. Henson BJ, Bhattacharjee S, O’Dee DM, Feingold E, Gollin SM. 2009. Decreased expression of miR-125b and miR-100 in oral cancer cells contributes to malignancy. Genes Chromosomes Cancer 48:569–582.

59. Wang Z, Wang J, Chen Z, Wang K, Shi L. 2018. MicroRNA-1-3p inhibits the proliferation and migration of oral squamous cell carcinoma cells by targeting DKK1. Biochem Cell Biol 96:355–364.

60. Wu Y-Y, Chen Y-L, Jao Y-C, Hsieh I-S, Chang K-C, Hong T-M. 2014. miR-320 regulates tumor angiogenesis driven by vascular endothelial cells in oral cancer by silencing neuropilin 1. Angiogenesis 17:247–260.

61. Schneider A, Victoria B, Lopez YN, Suchorska W, Barczak W, Sobecka A, Golusinski W, Masternak MM, Golusinski P. 2018. Tissue and serum microRNA profile of oral squamous cell carcinoma patients. Sci Rep 8:675.

62. Granda-Díaz R, Manterola L, Hermida-Prado F, Rodríguez R, Santos L, García-de-la-Fuente V, Fernández MT, Corte-Torres MD, Rodrigo JP, Álvarez-Teijeiro S, Lawrie CH, Garcia-Pedrero JM. 2023. Targeting oncogenic functions of miR-301a in head and neck squamous cell carcinoma by PI3K/PTEN and MEK/ERK pathways. Biomed Pharmacother 161:114512.

63. Liu J, Shi W, Wu C, Ju J, Jiang J. 2014. miR-181b as a key regulator of the oncogenic process and its clinical implications in cancer (Review). Biomed Rep 2:7–11.

64. Liu B, Chen W, Cao G, Dong Z, Xu J, Luo T, Zhang S. 2017. MicroRNA-27b inhibits cell proliferation in oral squamous cell carcinoma by targeting FZD7 and Wnt signaling pathway. Arch Oral Biol 83:92–96.

65. Zhang Y, Jin X, Wang J. 2019. miRL148a modulates the viability, migration and invasion of oral squamous cell carcinoma cells by regulating HLALG expression. Mol Med Rep 20:795–801.

66. Luo C, Zhang J, Zhang Y, Zhang X, Chen Y, Fan W. 2020. Low expression of miR-let-7a promotes cell growth and invasion through the regulation of c-Myc in oral squamous cell carcinoma. Cell Cycle 19:1983–1993.

67. Wang X, Li G-H. 2018. MicroRNA-16 functions as a tumor-suppressor gene in oral squamous cell carcinoma by targeting AKT3 and BCL2L2. J Cell Physiol 233:9447– 9457.

68. Fukumoto I, Hanazawa T, Kinoshita T, Kikkawa N, Koshizuka K, Goto Y, Nishikawa R, Chiyomaru T, Enokida H, Nakagawa M, Okamoto Y, Seki N. 2015. MicroRNA expression signature of oral squamous cell carcinoma: functional role of microRNA-26a/b in the modulation of novel cancer pathways. Br J Cancer 112:891–900.

69. Wu S, Li D, Han P, Li L, Zhao J, Zhang H, Zhou X, Li P, Mo Y. 2024. MicroRNAL374aL5p/ANLN axis promotes malignant progression of Oral squamous cell carcinoma. Nucleosides Nucleotides Nucleic Acids 1–16.

70. Fukumoto I, Koshizuka K, Hanazawa T, Kikkawa N, Matsushita R, Kurozumi A, Kato M, Okato A, Okamoto Y, Seki N. 2016. The tumor-suppressive microRNA-23b/27b cluster regulates the MET oncogene in oral squamous cell carcinoma. Int J Oncol 49:1119–1129.

71. Mitsueda R, Toda H, Shinden Y, Fukuda K, Yasudome R, Kato M, Kikkawa N, Ohtsuka T, Nakajo A, Seki N. 2023. Oncogenic Targets Regulated by Tumor-Suppressive miR-30c-1-3p and miR-30c-2-3p: TRIP13 Facilitates Cancer Cell Aggressiveness in Breast Cancer. Cancers (Basel) 15:4189.

72. Ayesha M, Majid A, Zhao D, Greenaway FT, Yan N, Liu Q, Liu S, Sun M-Z. 2022. MiR-4521 plays a tumor repressive role in growth and metastasis of hepatocarcinoma cells by suppressing phosphorylation of FAK/AKT pathway via targeting FAM129A. J Adv Res 36:147–161.

73. Sun B, Cong D, Chen K, Bai Y, Li J. 2021. Prognostic value of microRNA-4521 in non-small cell lung cancer and its regulatory effect on tumor progression. Open Med (Wars) 16:1150–1159.

74. Venkatesh T, Nagashri MN, Swamy SS, Mohiyuddin SMA, Gopinath KS, Kumar A. 2013. Primary microcephaly gene MCPH1 shows signatures of tumor suppressors and is regulated by miR-27a in oral squamous cell carcinoma. PLoS One 8:e54643.

75. Tseng H-H, Tseng Y-K, You J-J, Kang B-H, Wang T-H, Yang C-M, Chen H-C, Liou H-H, Liu P-F, Ger L-P, Tsai K-W. 2017. Next-generation Sequencing for microRNA Profiling: MicroRNA-21-3p Promotes Oral Cancer Metastasis. Anticancer Res 37:1059–1066.

76. Chen Q, Wang W, Wang Y. 2022. MiR-222 regulates the progression of oral squamous cell carcinoma by targeting CDKN1B. Am J Transl Res 14:5215–5227.

77. Komatsu S, Ichikawa D, Kawaguchi T, Takeshita H, Miyamae M, Ohashi T, Okajima W, Imamura T, Kiuchi J, Arita T, Konishi H, Shiozaki A, Fujiwara H, Okamoto K, Otsuji E. 2016. Plasma microRNA profiles: identification of miR-23a as a novel biomarker for chemoresistance in esophageal squamous cell carcinoma. Oncotarget 7:62034–62048.

78. Guo J, Wen N, Yang S, Guan X, Cang S. 2018. MiR-92a regulates oral squamous cell carcinoma (OSCC) cell growth by targeting FOXP1 expression. Biomed Pharmacother 104:77–86.

79. Tripathi S, Sharma Y, Kumar D. 2025. Unveiling the link between chronic inflammation and cancer. Metabol Open 25:100347.

80. Fernandes Q, Inchakalody VP, Bedhiafi T, Mestiri S, Taib N, Uddin S, Merhi M, Dermime S. 2024. Chronic inflammation and cancer; the two sides of a coin. Life Sci 338:122390.

81. Greten FR, Grivennikov SI. 2019. Inflammation and Cancer: Triggers, Mechanisms, and Consequences. Immunity 51:27–41.

82. Miyajima C, Inoue Y, Hayashi H. 2015. Pseudokinase tribbles 1 (TRB1) negatively regulates tumor-suppressor activity of p53 through p53 deacetylation. Biol Pharm Bull 38:618–624.

83. Moyes DL, Shen C, Murciano C, Runglall M, Richardson JP, Arno M, Aldecoa-Otalora E, Naglik JR. 2014. Protection against epithelial damage during Candida albicans infection is mediated by PI3K/Akt and mammalian target of rapamycin signaling. J Infect Dis 209:1816–1826.

84. Naglik JR, Richardson JP, Moyes DL. 2014. Candida albicans pathogenicity and epithelial immunity. PLoS Pathog 10:e1004257.

85. Pellon A, Sadeghi Nasab SD, Moyes DL. 2020. New Insights in Candida albicans Innate Immunity at the Mucosa: Toxins, Epithelium, Metabolism, and Beyond. Front Cell Infect Microbiol 10:81.

86. Swidergall M, Solis NV, Lionakis MS, Filler SG. 2018. EphA2 is an epithelial cell pattern recognition receptor for fungal β-glucans. Nat Microbiol 3:53–61.

87. Mostefaoui Y, Claveau I, Rouabhia M. 2004. In vitro analyses of tissue structure and interleukin-1beta expression and production by human oral mucosa in response to Candida albicans infections. Cytokine 25:162–171.

88. Aparicio-Fernandez L, Cazalis-Bereicua N, Areitio M, et al. 2025. Candida albicans enhances melanoma cell aggressiveness through p38-MAPK and HIF-1α pathways and metabolic reprogramming 10.1101/2025.01.17.633543.

89. Kakade P, Burgueno JF, Sircaik S, Ponde N, Li J, Ene IV, Kim J, Liang S-H, Yunker R, Akiba Y, Vaishnava S, Kaunitz JD, Way SS, Koh AY, Gaffen S, Abreu MT, Bennett RJ. 2024. An IL-17-DUOX2 axis controls gastrointestinal colonization by Candida albicans. bioRxiv 2024.08.16.608271.

90. Merayo-Lloves J, Quirós LM, Martin C, Alcalde I, Iturbe VL, Ordiales-Trabanco H. 2021. Role in adhesion and changes in cell surface proteoglycan expression in Candida keratitis. iovs 62:1957.

91. Dweep H, Gretz N. 2015. miRWalk2.0: a comprehensive atlas of microRNA-target interactions. Nat Methods 12:697.

