## Supplementary figures and images for "Early *Candida*–Oral Tumor Interactions Suggest miRNA-Mediated Regulation of Inflammatory and Tumor-associated processes"

### FIG S1

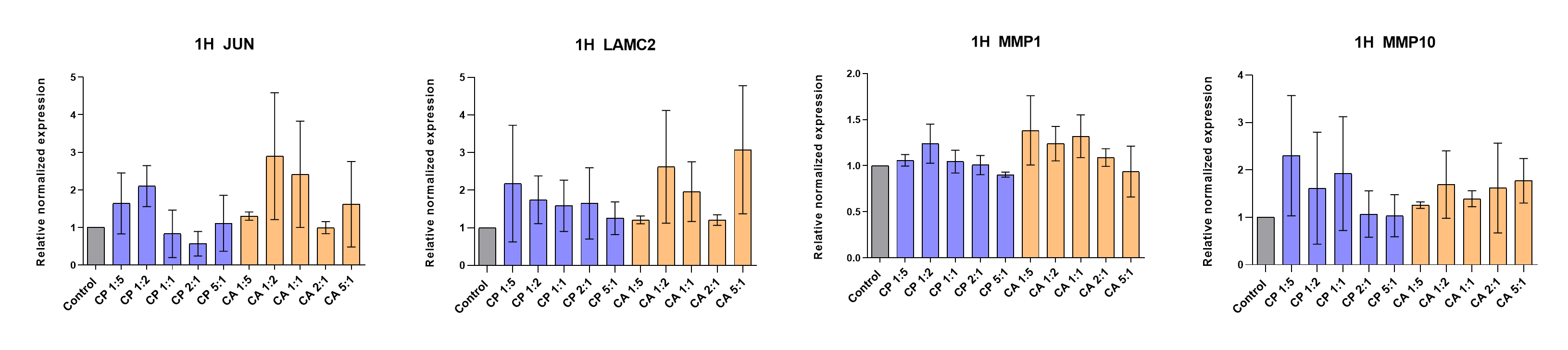

### FIG S2A-C

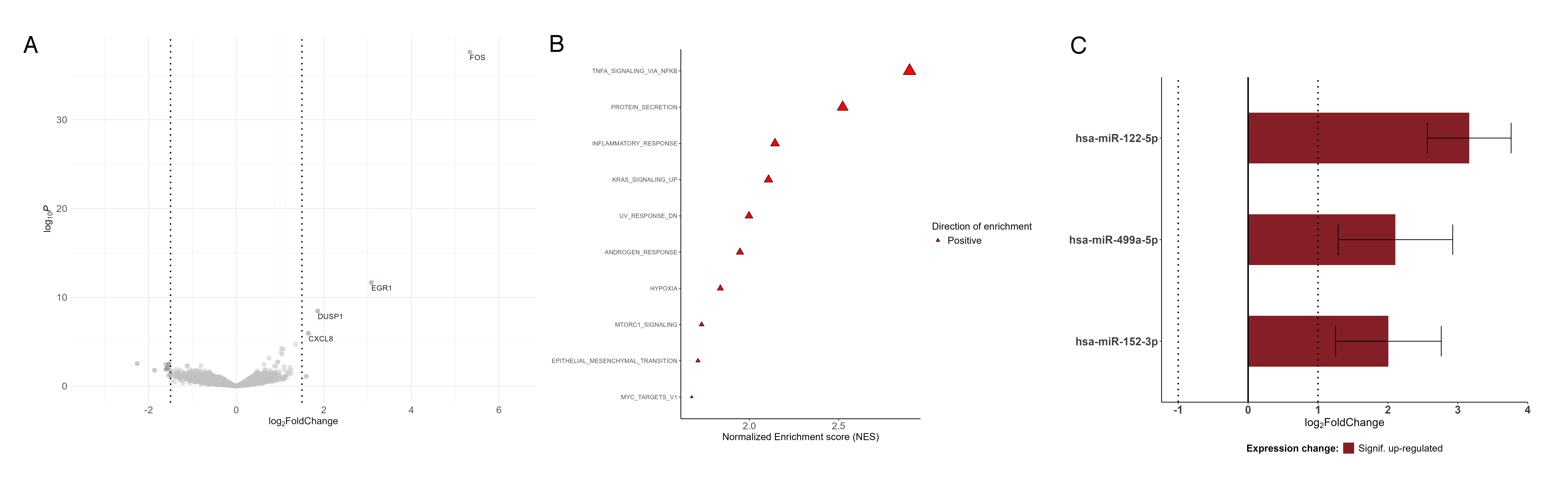

### FIG S3

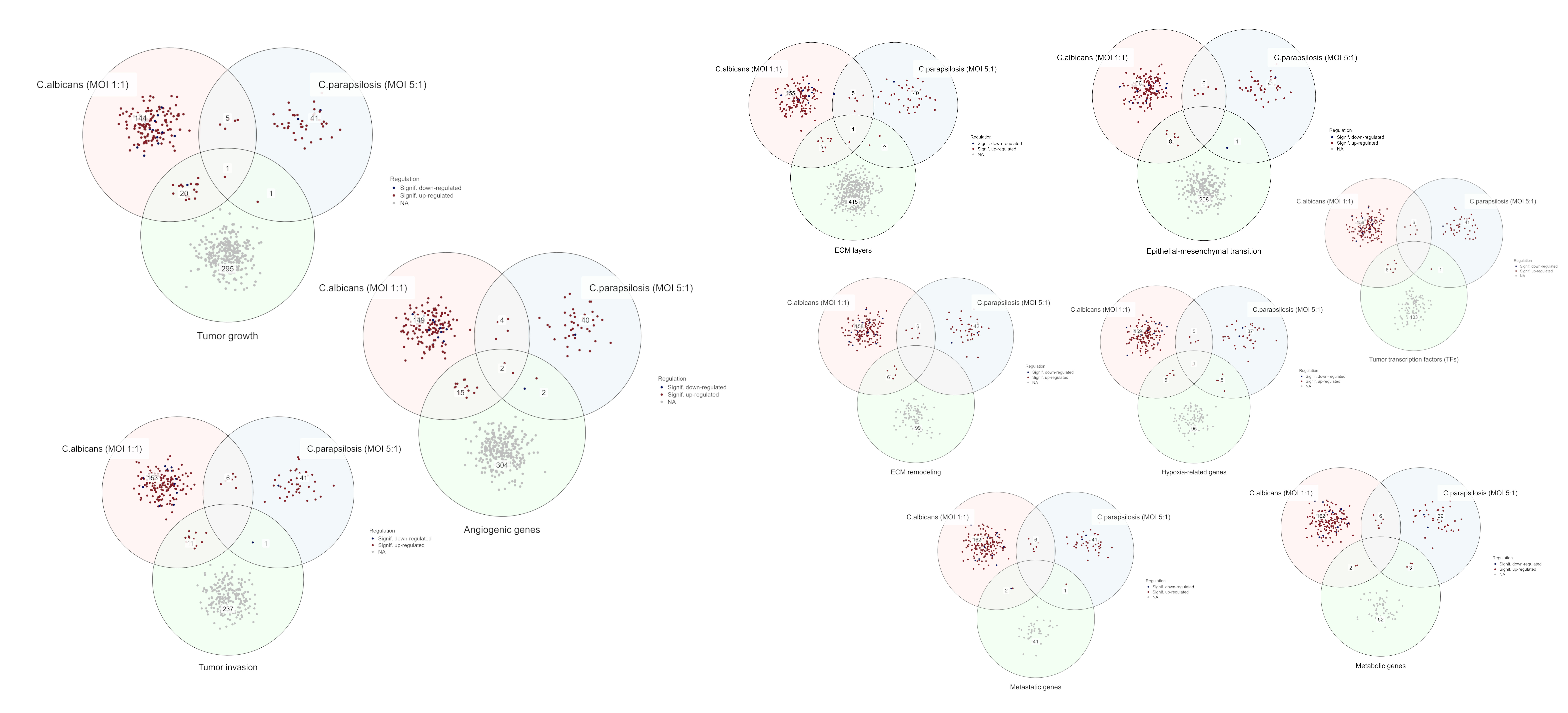

### FIG S4A-B

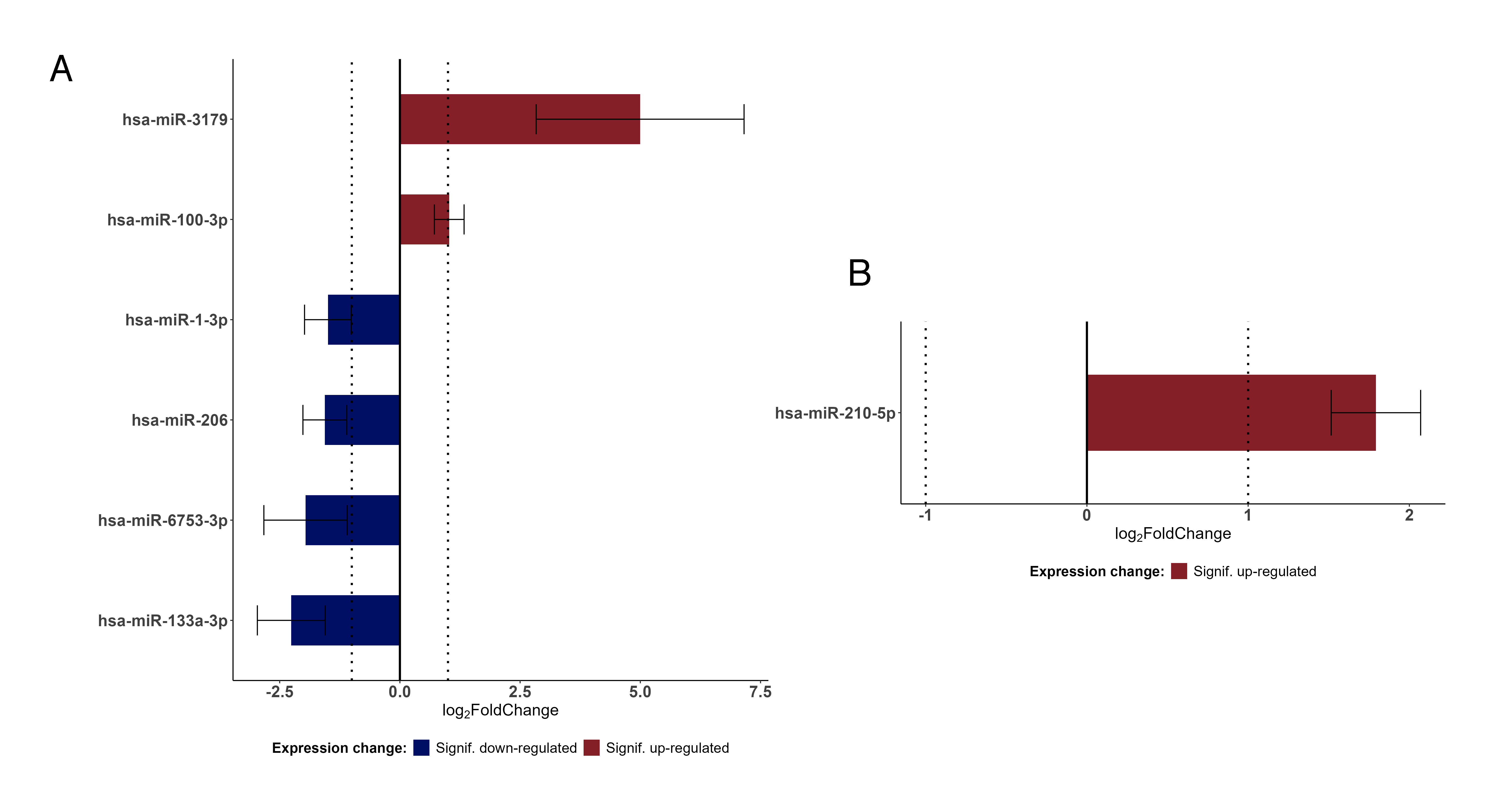
